## Supplementary Material for "Differential elasticity in lineage segregation of embryonic stem cells"

**Cell culture.** mCherry-Hhex as well as constitutive expressing H2B-Venus labeled mESCs were described previously [1]. Cells were grown in culture flasks (BD Falcon) coated with 0.1 % gelatin (Millipore) in a cell culture incubator (Thermo Scientific) at 37° C and 5 % $CO_2$ using Glasgow minimum essential medium (GMEM, Sigma, Germany) with 10 % fetal bovine serum (FBS) and leukemia inhibitor factor (LIF). The medium was furthermore enriched with 5.5 ml non-essential amino acids (Gibco), 5.5 ml L-glutamine (Gibco), 5.5 ml sodium pyruvate (Gibco) and 0.1 mM mercaptoethanol (Sigma, Germany).
For experiments, cells were plated on a 0.1 % gelatin-coated coverslip (24 x 60 mm, #1.5, VWR) and incubated for 12-24 hours in the incubator at 5 % $CO_2$ and 37° C allowing them to adhere. To create a perfusion chamber two stripes of vacuum grease were applied to the long sides of the cell coated coverslips. A second clean coverslip (18 x 18 mm, #1, Menzel-Gläser) was placed on top of the vacuum grease stripes thereby creating a chamber. This chamber was filled with culture medium and the two open sides were sealed with vacuum grease. The samples were prepared and investigated at room temperature ($\sim$ 24° C). For actin depolymerization experiments, cells were treated with 1 $\mu$M latrunculin B (Sigma, Germany) for 1 hour prior to measurements.

**Optical tweezers tracking.** The positions of endogenous lipid granules (in the lateral plane) were tracked using optical tweezers. The optical tweezers were integrated in a confocal microscope (TCS SP5, Leica Microsystems GmbH, Germany) by tightly focusing a near infrared (Nd:YVO4, 5 W Spectra Physics BL106C, $\lambda = 1064$ nm, TEM$\infty$) through the microscope objective (HCX PL APO, 63x, NA=1.2, COR R CS). After passing the sample, the laser light was collected by a condenser and imaged onto a quadrant photodiode, details on the equipment can be found in Ref. [2, 3]. A custom-written LabVIEW program (LabVIEW, National Instruments) was used to acquire a time series of the granules' position in the lateral directions. The measurements were conducted by focusing the optical trap on individual granules within the cytoplasm or the nucleus of EPI- or PrE-primed cells, which were kept at room temperature. The trapping laser was operated at a laser power of 140 mW and each measurement lasted 3 seconds. At this irradiation power and duration, the laser induced heating is less than 1° C and there is no observable physiological damage [4–6].

**Image analysis.** The confocal mode of the microscope was used to determine the primed state of cells by measuring its mCherry expression under the Hhex promoter. LAS AF software (Leica Microsystems GmbH, Germany) was used to determine the average fluorescence mCherry-Hhex intensities of a region of interest inside an individual cell. Cells were categorized into three different groups: (i) Cells exhibiting low mCherry-Hhex intensities indistinguishable from auto-fluorescence were considered EPI-primed cells. (ii) Cells with medium mCherry-Hhex intensities were considered PrE-primed. (iii) Spontaneous differentiation into the PrE-lineage can occur in cell culture. These fully differentiated cells were recognizable by their high Hhex expression and different morphology and were avoided. After determining their priming, the cells were imaged by bright field microscopy, thus allowing for identifying lipid granules inside the cells, and the laser trap was focused on one such granule.

**Time series analysis.** The time series of the granule's position was Fourier transformed to $\tilde{x}(f)$, where $f$ denotes the frequency. From this the power spectrum was calculated as

$$P(f) = \frac{|\tilde{x}(f)|}{T_{\text{meas}}}, \quad (1)$$

where $T_{\text{meas}}$ is the total measurement time. A MATLAB (MathWorks, Inc., USA) program adapted from [3, 7] was used for calculation of $P(f)$ and for fitting $P(f) \propto f^{-(1+\alpha)}$ to the data. The frequency below which an optically trapped tracer feels the restoring potential of the optical trap is denoted the corner frequency, $f_c$. All power spectra were fitted in the frequency interval 200 Hz-3000 Hz, which was well beyond $f_c$ ($f_c \ll 200$ Hz) and below the 3 dB filtering frequency of the quadrant

* These authors contributed equally to this work

photodiode [8].

**Statistical analysis.** Statistical analysis was performed using Prism software (GraphPad, version 9.4.0). $p$-values were obtained using Welch's $t$-tests. Graphs show symbols describing $p$-values: *: $p < 0.05$; **: $p < 0.01$; ***: $p < 0.001$, ****: $p < 0.0001$. Black line in scatter plots denotes the median. Data in Fig 2a) stems from 100 cells (untreated: 56 EPI and 44 PrE-primed) and in Fig 2b) from 53 cells (LatB treated: 28 EPI and 25 PrE-primed).

**Calculation of the shear modulus.** Viscoelastic properties of cells or materials can be quantified with the frequency dependent complex shear modulus, $G(f) = G'(f) + iG''(f)$, where $G'(f)$ reports the elastic storage response and $G''(f)$ reports the viscous loss response. The loss tangent, the ratio between the loss and storage moduli $G''/G'$, reflects the degree of solid- or fluid-like behavior of the cell [9]. The Fourier transform of the displacement of a particle in a viscoelastic medium, $\tilde{x}(f)$, is linearly related to the stochastic force from the surrounding medium, $F(f)$, with the response function, $\gamma(f)$ [10, 11]:

$$\tilde{x}(f) = \gamma(f)F(f). \tag{2}$$

The Stoke-Einstein relates the shear modulus to the complex response function, $\gamma(f)$:

$$G(f) = \frac{1}{6\pi r\gamma(f)}, \tag{3}$$

where $r$ is the radius of the particle in the trap. Based on the fluctuation-dissipation theorem, which is valid under the current experimental conditions [12], the imaginary part of the response function, $\gamma''(f)$, can be extracted from power spectral density, $P(f)$:

$$\gamma''(f) = \frac{\pi f}{2K_{\mathrm{B}}\mathrm{T}}P(f), \tag{4}$$

where $K_{\mathrm{B}}$ is the Boltzmann constant and T is the temperature. The real part of $\gamma(f)$ can be calculated numerically using a Kramers–Kronig transformation:

$$\gamma'(f) = \frac{2}{\pi}P\int_{f''=0}^{f''=\infty} df'' \frac{f''\gamma'(f'')}{f''^2 - f^2}. \tag{5}$$

Finally, substituting Eq. 5 and Eq. 4 into Eq. 3 gives the storage and loss moduli:

$$G'(f) = \frac{1}{6\pi r}\frac{\gamma'(f)}{\gamma'(f)^2 + \gamma''(f)^2}, \tag{6}$$

$$G''(f) = \frac{-1}{6\pi r}\frac{\gamma''(f)}{\gamma'(f)^2 + \gamma''(f)^2}. \tag{7}$$

For a particle in the trap, $G'(f)$ comprises a contribution from the elastic modulus of the trap $G'_{\mathrm{trap}}(f)$, and the surrounding medium. Therefore, in our calculations we subtracted the elastic modulus of the trap from results in Fig. S3. To estimate the elastic modulus of the trap, we trapped a bead in water with the same laser intensity we used for trapping granules inside cells and found the spring constant characterizing the harmonic optical trapping potential. As the elastic modulus of water is close to zero, the total elastic modulus only contains the contribution from the trap (Fig. S3). By extracting the shear modulus inside EPI- and PrE-primed cells we found that in EPI-primed cells, the loss modulus dominates over the storage modulus in the whole measured frequency range (1-500 Hz). Hence, EPI-primed cells show fluid-like behavior ($G''/G' > 1$), whereas PrE cells exhibit more solid-like behavior up to 15 Hz ($G''/G' < 1$), where the loss modulus crosses over the storage modulus (Fig. 2c,d). Comparing our results to reported values for other types of cells, we find that the storage modulus of PrE-primed cells (215-1500 Pa, 1-100 Hz) is comparable with MCF7 cells (∼400-1000 Pa), MDA-MB-231 cells (∼700-2000 Pa) [13], and MDCK-II cells (∼300-1000 Pa) [14]. In contrast, the storage modulus of EPI-primed cells, 20-360 Pa, is significantly lower than for PrE-primed and other cells reported in literature. This indicates that the EPI-primed cells are more viscous than most other cells whereas, PrE-primed cells (in our study) and other reported cells [13, 14] show a predominantly solid-like behavior.

**Computational model.** To explore the effect of differential elasticity on cell segregation we employ a cell-based model, where cell deformation, motility, and cell-cell interactions are captured at the individual cell level [15–20]. This model is recently proven successful in describing the emergence of orientational order and topological defects in epithelial tissues [21, 22] and has been quantitatively compared with the results of sustained oscillations in confined human skin cells [19], as well as with experimental results on mechanisms of force generation in epithelial and mesenchymal cells [23]. We therefore refer the reader to [17, 19, 21] for details of the implementation of the model, and provide a brief description of the framework here. Each cell is described by a single phase-field $\Phi$ where $\Phi_i \simeq 1$ denotes the interior of a cell and $\Phi_i \simeq 0$ its exterior. The boundary of each cell is described implicitly and is defined to lie at $\Phi_i \simeq 1/2$. The dynamics of the phase-fields $\Phi$ are given by

$$\partial_t \Phi_i + \boldsymbol{v}_i \cdot \nabla \Phi_i = -\frac{\delta \mathcal{F}}{\delta \Phi_i}, \tag{8}$$

where $\mathcal{F}$ is the free energy of the system and the advection term is the source of the non-equilibrium properties of the model. Our model effectively describes cells as active deformable droplets. The free energy $\mathcal{F}$ controls the physical properties of the cells as well as their interactions. We decompose it as

$$\mathcal{F} = \underbrace{\mathcal{F}_{\text{CH}} + \mathcal{F}_{\text{area}}}_{F_{\text{cell}}} + \underbrace{\mathcal{F}_{\text{rep}} + \mathcal{F}_{\text{adh}}}_{F_{\text{int}}}, \tag{9}$$

where $\mathcal{F}_{\text{CH}}$ is the double-well free energy responsible for describing the diffusive interface, $\mathcal{F}_{\text{area}}$ implements a constraint for area conservation, while $\mathcal{F}_{\text{rep}}$ and $\mathcal{F}_{\text{adh}}$ give rise to repulsion and adhesion between cells, respectively:

$$\mathcal{F}_{\text{area}} = \sum_i \int d^2\boldsymbol{x}\mu \left(1 - \frac{\int dx \Phi_i^2}{\pi R^2}\right)^2, \tag{10}$$

$$\mathcal{F}_{\text{rep}} = \sum_i \sum_{i \neq j} \frac{30\kappa}{\lambda^2} \Phi_i^2 \Phi_j^2, \tag{11}$$

$$\mathcal{F}_{\text{adh}} = \sum_i \sum_{i \neq j} \frac{30\omega}{\lambda^2} (\nabla\Phi_i)^2 (\nabla\Phi_j)^2, \tag{12}$$

where $\kappa$ and $\omega$ set the strength of the repulsion and adhesion between the cells, respectively, and $\mu$ sets the compressibility of the cell. The contribution $\mathcal{F}_{\text{CH}}$ in Eq. 9 controls the stiffness of the cells and is given by

$$\mathcal{F}_{\text{CH}} = \sum_i \int d^2\boldsymbol{x} \frac{\gamma_i}{\lambda} \left(4\Phi_i^2(1-\Phi_i)^2 + \lambda^2(\nabla\Phi_i)^2\right), \tag{13}$$

where $\gamma_i$ is the cell stiffness and $\lambda$ defines the length-scale of the diffusive interface, see [19] for more details. In our simulations we set $\gamma_i$ to different values for the two cell types with smaller $\gamma_i$ representing the fluid-like, less stiff, cells and larger $\gamma_i$ corresponding to solid-like, stiffer, cells. As it was recently demonstrated a measure of solid- versus fluid-like behavior can be obtained by calculating the neighbor exchange between the cells in a monolayer [24]. We quantified this solid- versus fluid-like behavior by measuring the rate of neighbor exchange between the cells in monolayers with low and high $\gamma_i$ as shown in Fig. S6. As discussed in the main text, an essential factor for the elastic segregation is the ability of cells to generate active stresses that puts the system out of thermodynamic equilibrium. Such active stresses are mainly produced by acto-myosin contractility of the cell and play an important role in cell motility [25–27]. In our formulation, the non-equilibrium effects of active stress generation and cell motility enter through an overdamped velocity $\boldsymbol{v}_i$ in Eq. 8. The force balance is used to determine the velocity in Eq. 8. Balancing the friction force acting on the substrate with the sum of all intercellular forces and active forces, we write

$$\xi \boldsymbol{v}_i = \boldsymbol{F}_i^{\text{int}} + \boldsymbol{F}_i^{\text{active}}, \tag{14}$$

where $\xi$ is a friction coefficient. The intercellular and active forces are defined in terms of the corresponding stresses as [21]:

$$\boldsymbol{F}_i^{\text{int}} = \int d^2\boldsymbol{x}\ \nabla \cdot \boldsymbol{\sigma}^{\text{int}} = -\int d^2\boldsymbol{x}\ \boldsymbol{\sigma}^{\text{int}} \cdot \nabla\Phi_i, \tag{15}$$

$$\boldsymbol{F}_i^{\text{active}} = \int d^2\boldsymbol{x}\ \nabla \cdot \boldsymbol{\sigma}^{\text{active}} = -\int d^2\boldsymbol{x}\ \boldsymbol{\sigma}^{\text{active}} \cdot \nabla\Phi_i. \tag{16}$$

The intercellular stress $\boldsymbol{\sigma}^{\text{active}}$ accounts for the passive stresses arising due to cell-cell interactions and is determined from the free energy $\mathcal{F}$:

$$\boldsymbol{\sigma}^{\text{int}} = -\sum_i \frac{\delta\mathcal{F}}{\delta\Phi_i}\mathbf{I}, \tag{17}$$

where $\mathbf{I}$ is the identity tensor, while the active stress governs the cell deformation:

$$\boldsymbol{\sigma}^{\text{active}} = -\xi \sum_j \Phi_j \mathbf{S}_j, \tag{18}$$

where $\xi$ is the strength of active stress and $\mathbf{S}_i = -\int d^2\boldsymbol{x}(\nabla\Phi_i)^{\text{T}}\nabla\Phi_i$ is the deformation tensor for each cell. As shown previously [21], and compared with the mechanisms of force generation in epithelial and mesenchymal cell layers [23], this form of active stress results in the force-dipole generation by each cell which represents the force-dipoles exerted by individual cells on their underlying substrates due to their internal acto-myosin contractility. Therefore, the intercellular and active forces are written as:

$$\boldsymbol{F}_i^{\text{int}} = \int d^2\boldsymbol{x}\ (\frac{\delta\mathcal{F}}{\delta\Phi_i}\mathbf{I}), \tag{19}$$

$$\boldsymbol{F}_i^{\text{active}} = \xi \int d^2\boldsymbol{x} \left(\sum_j \Phi_j \mathbf{S}_j\right) \cdot \nabla\Phi_i. \tag{20}$$

These expressions can be easily understood by realizing that the gradient $\nabla\Phi_i$ is non-zero only at the cell interface and points towards its exterior. Hence, they amount in computing to the total force on one cell's interface. We simulate equations (4-7) using a finite-difference scheme on a square lattice with a predictor-corrector step. Thanks to the algorithm described in [21], we consider only nodes that are included in a subdomain associated with a cell and are able to achieve linear computational and memory complexity with respect to the number of cells. We implemented our algorithm in C++ and parallelized it to multi-core architectures using OpenMP and to GPU using Cuda. We have observed that GPU acceleration improves this baseline by about a factor of 100 using a GeForce GTX 1080 Ti. All the double sums appearing in $\frac{\delta\mathcal{F}}{\delta\Phi_i}$ can be dealt with efficiently by storing global fields such as $\Phi_i$ at every point of the domain. Because the total force involves integrals over the whole domain we use a 4 simple two-pass algorithm where the phase-fields $\Phi_i$ are updated one after another. Unless otherwise stated, simulation parameters are $\gamma = 0.01$, $\mu = 3$, $\lambda = 2.5$, $\kappa = 0.1$, $R = 8$, $\xi = 2$ and we use a subdomain size of 25×25 lattice sites. All simulation results are reported in terms of the dimensionless numbers $(\gamma_A - \gamma_B)/\gamma_B$ and $\zeta R/\bar{\gamma}$, where the former characterizes the elasticity difference and the latter ratio of active stress to the average elasticity $\bar{\gamma} = (\gamma_A + \gamma_B)/2$.

| Category | $N_{untreated}$ | $\alpha$ *untreated* | $N_{LatB}$ | $\alpha$ *LatB* |
|---|---|---|---|---|
| EPI | 642 | 0.49±0.13 | 450 | 0.52±0.09 |
| $EPI_{nucleus}$ | 327 | 0.47±0.13 | 225 | 0.49±0.09 |
| $EPI_{cytoplasm}$ | 315 | 0.51±0.12 | 225 | 0.56±0.07 |
| PrE | 759 | 0.42±0.17 | 465 | 0.55±0.09 |
| $PrE_{nucleus}$ | 372 | 0.39±0.16 | 210 | 0.55±0.08 |
| $PrE_{cytoplasm}$ | 387 | 0.45±0.17 | 255 | 0.56±0.09 |

TABLE I. **Information about the average value of scaling exponents, $\alpha$, as well as the number of experiments, $N$, in each category.** Average values of $\alpha$ for all measurements on EPI- and PrE-primed cells for granules irrespective of location within the cell and split up to granules inside the nucleus and inside the cytoplasm, respectively. Columns highlighted in gray show data from normal (untreated) cells and columns highlighted in blue show data from cells treated with Latrunculin B (LatB) which depolymerizes actin.

| treatment | location | comparison | N (granules) | *p*-values | type of test |
|---|---|---|---|---|---|
| **untreated** | pooled (nucleus and cytoplasm) | EPI vs PrE | 642 and 759 | < 0.0001 | Welch's *t*-test |
| | nucleus | EPI vs PrE | 327 vs 372 | < 0.0001 | Welch's *t*-test |
| | cytoplasm | EPI vs PrE | 315 vs 387 | < 0.0001 | Welch's *t*-test |
| **LatB treated** | pooled (nucleus and cytoplasm) | LatB EPI vs LatB PrE | 450 vs 465 | | |
| | nucleus | EPI vs LatB EPI | 327 vs 225 | < 0.0001 | Welch's *t*-test |
| | | PrE vs LatB PrE | 372 vs 210 | < 0.0001 | Welch's *t*-test |
| | cytoplasm | EPI vs LatB EPI | 315 vs 225 | < 0.0001 | Welch's *t*-test |
| | | PrE vs LatB PrE | 387 vs 255 | < 0.0001 | Welch's *t*-test |

TABLE II. $p$-values from t-tests performed on pairs of distributions of experimentally obtained scaling exponents, $\alpha$ for normal, untreated cells (highlighted in gray), and cells treated with Latrunculin B (LatB) (highlighted in blue).

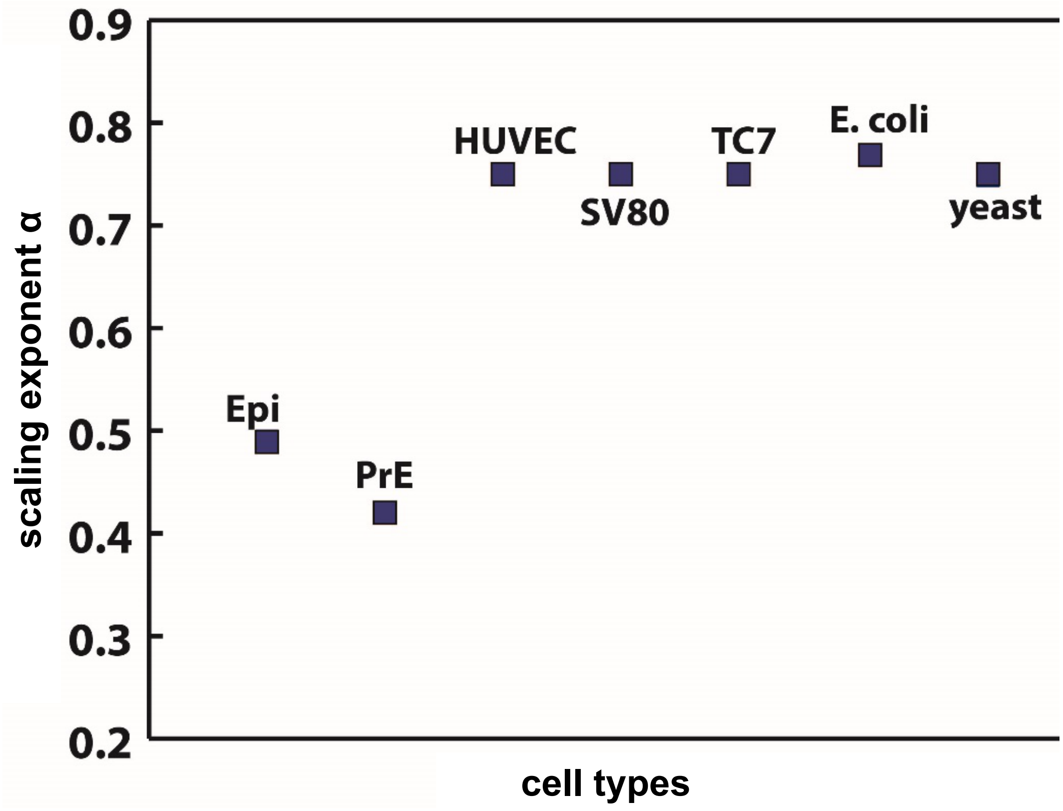


FIG. 1. **Comparison of the scaling exponents characterizing the intracellular micro-rheology of different cell types.** The ESCs (EPI- and PrE-primed) are in this investigation found to have scaling exponents that are significantly lower than any other cell type probed by using passively diffusing tracers, indicating that the measured cells in this study are significantly more elastic. The numbers depicted are taken from the following references: HUVEC [28], yeast [29], *E. coli* [30], TC7 [15], and SV80 [31]. If the tracers are carried by molecular motors, they typically perform a super-diffusive motion characterized by $\alpha > 1$.

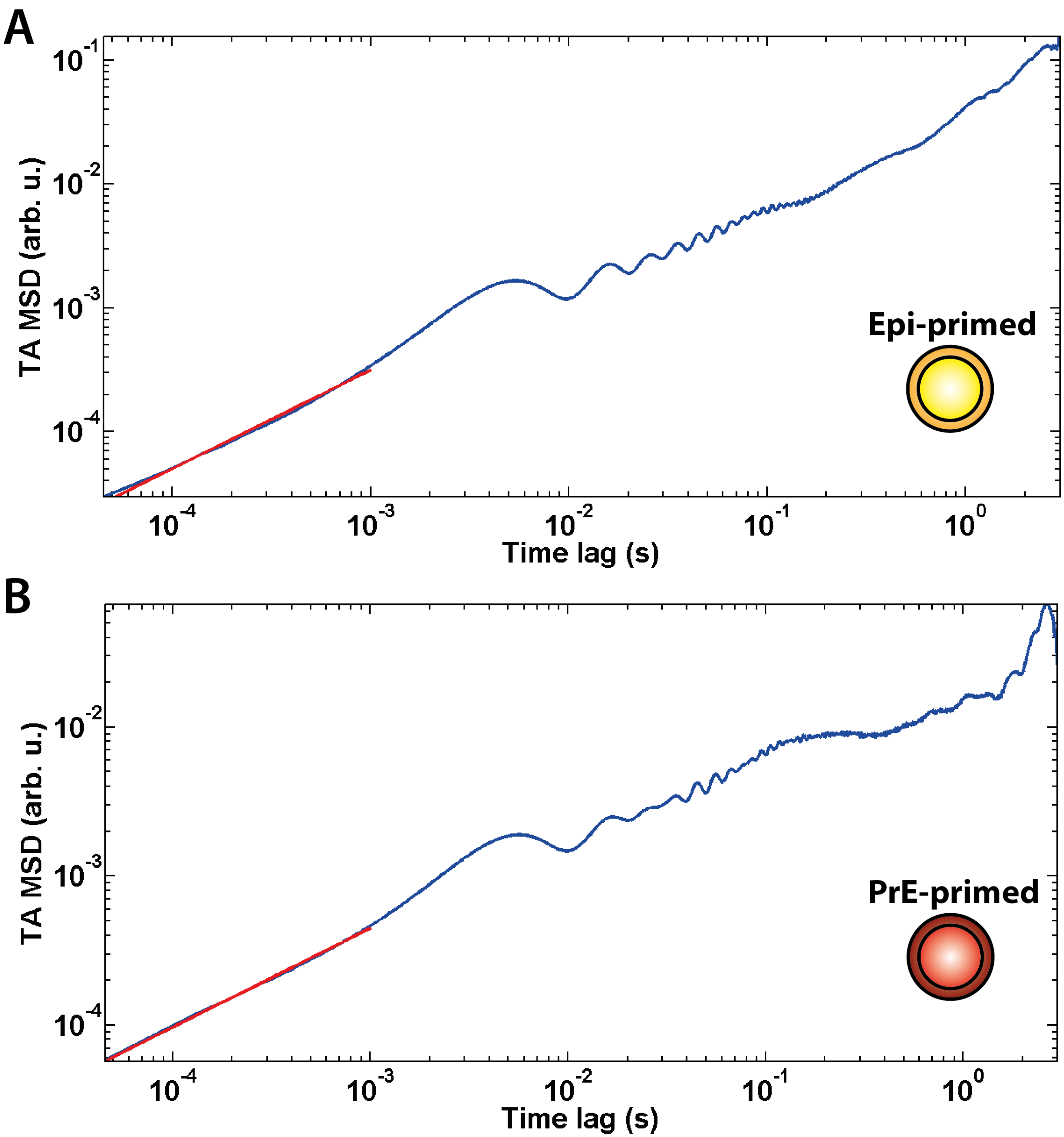


FIG. 2. Time averaged mean squared displacement (TA MSD) calculated from the trajectories of single lipid granules within A) an EPI-primed cell and B) a PrE-primed cell as a function of time lag. The TA MSDs were fitted at short time scales, up to 0.001 s, and the scaling exponents, , were extracted, giving $\alpha_{\mathrm{EPI}} = 0.80$ and $\alpha_{\mathrm{PrR}} = 0.66$ for the A) EPI-primed and the B) PrE-primed cell, respectively. At longer time-lags the TA MSD display oscillatory noise due to low-frequency electrical noise and mechanical vibrations.

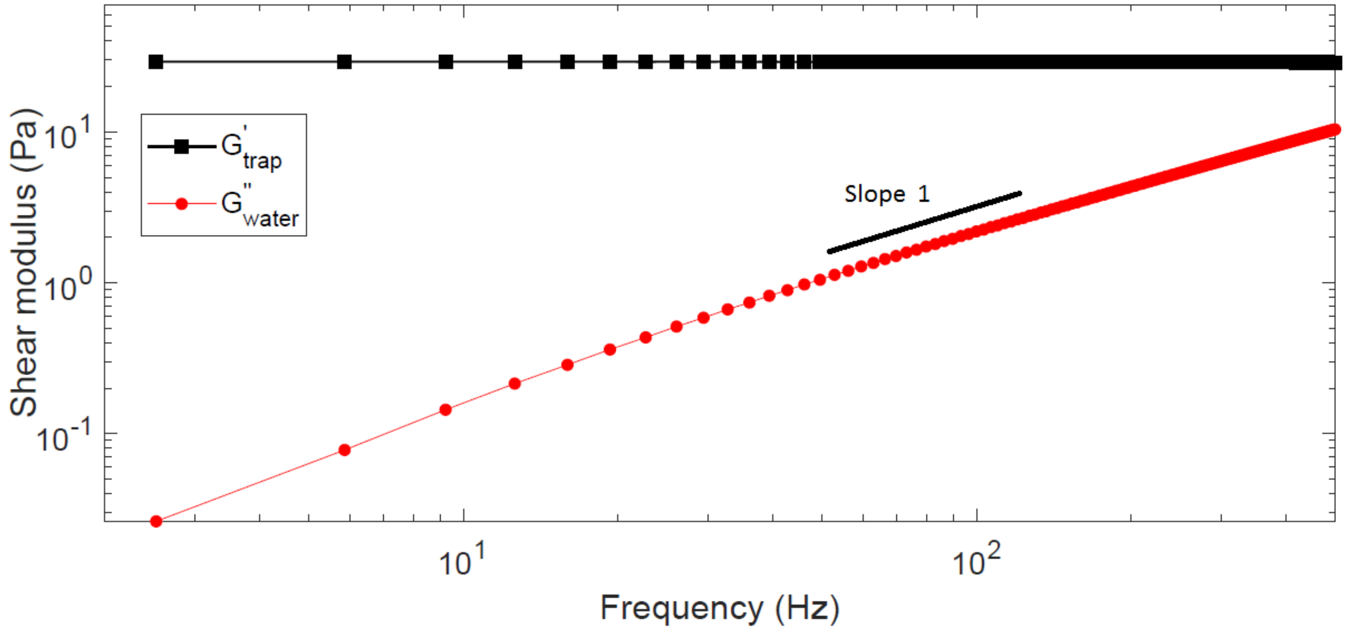


FIG. 3. **Elastic storage modulus of the optical trap.** The storage modulus of the trap (black squares) is frequency independent about 29 Pa in our study for a trapped bead in water, a purely viscous liquid. The loss modulus (red circles) is linearly increasing with frequency with an exponent of $\alpha = 1$ as expected for purely viscous liquid.

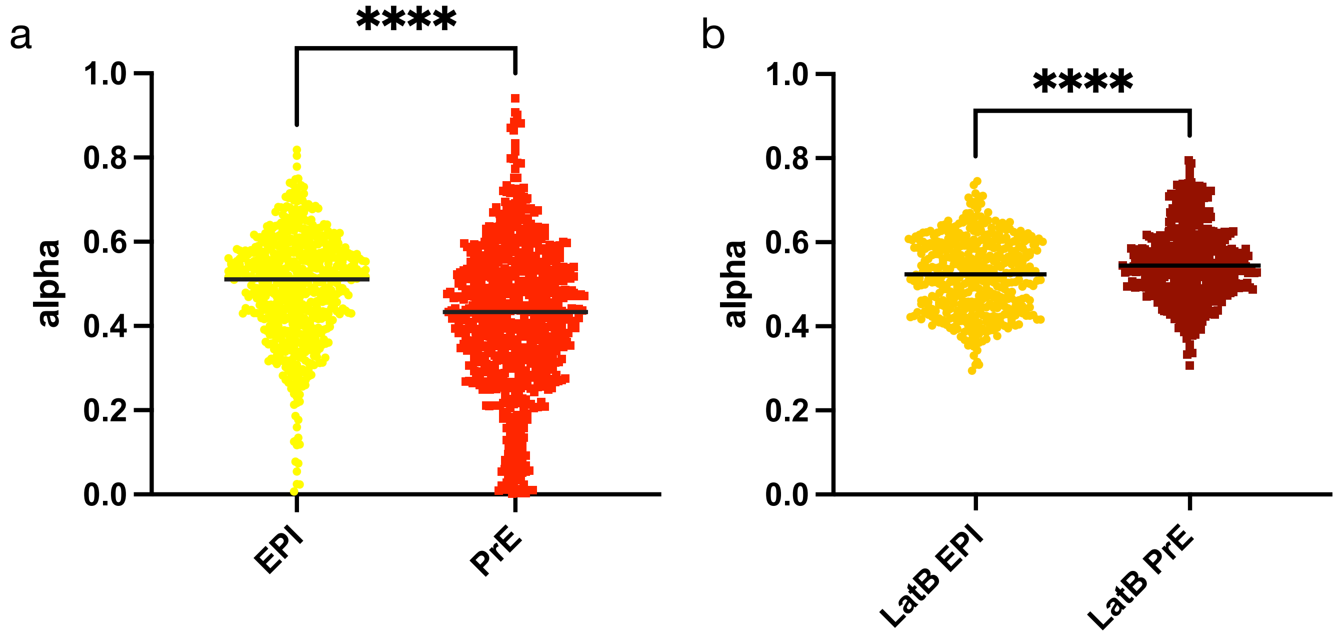


FIG. 4. Scatter plots of pooled scaling exponents $\alpha$ from a) untreated cells (EPI- and PrE-primed) and b) LatB treated cells (EPI- and PrE-primed). $p$-values and means are listed in Table S1 and S2. Black lines denote the median.

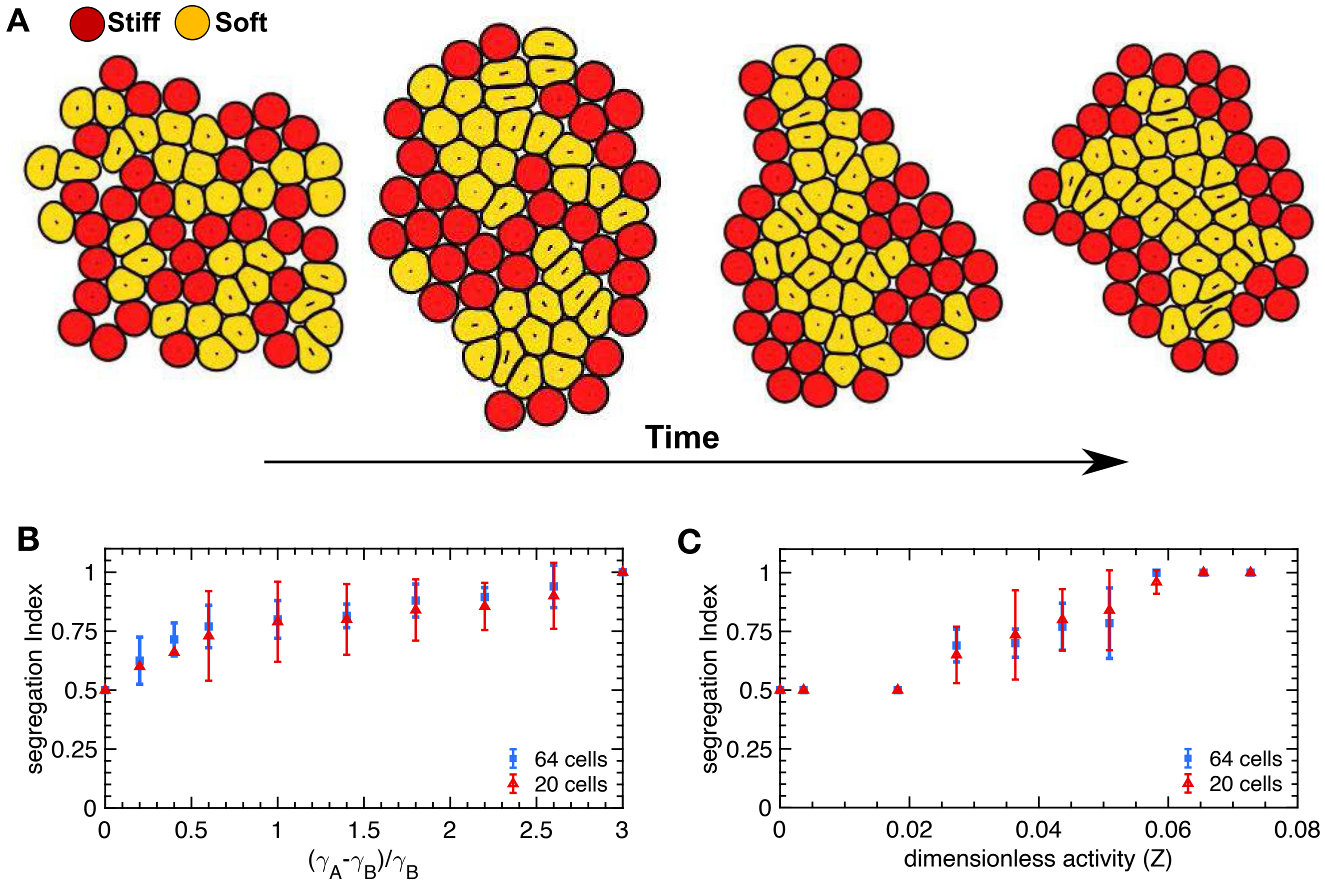


FIG. 5. **The effect of cell number on active viscoelastic segregation.** A) Time series snapshots of the segregation of a 64-cell aggregate of soft (yellow) and stiff (red) cells. B) Quantification of the effect of cell number on the cell segregation as a function of the normalized elasticity difference $(\gamma_A - \gamma_B)/\gamma_B$, where $\gamma_A$ denotes the elasticity of the stiff cells and $\gamma_B$ is the elasticity of the soft cells. C) Quantification of the cell number effect on the cell segregation as a function of the strength of active stresses generated by the cells. The segregation index $SI = \langle N_A/(N_A + N_B)\rangle$ is defined such that $SI = 1/2$ when the outer layer is a uniform mixture of stiff and soft cells (indicated by the dashed red line), $SI = 1$ when the outer layer is occupied solely by stiff cells, and $SI = -1$ if only soft cells occupy the outer layer. Error bars indicate the variance computed from averaging over 5 different simulations with random initial positioning of the cells.

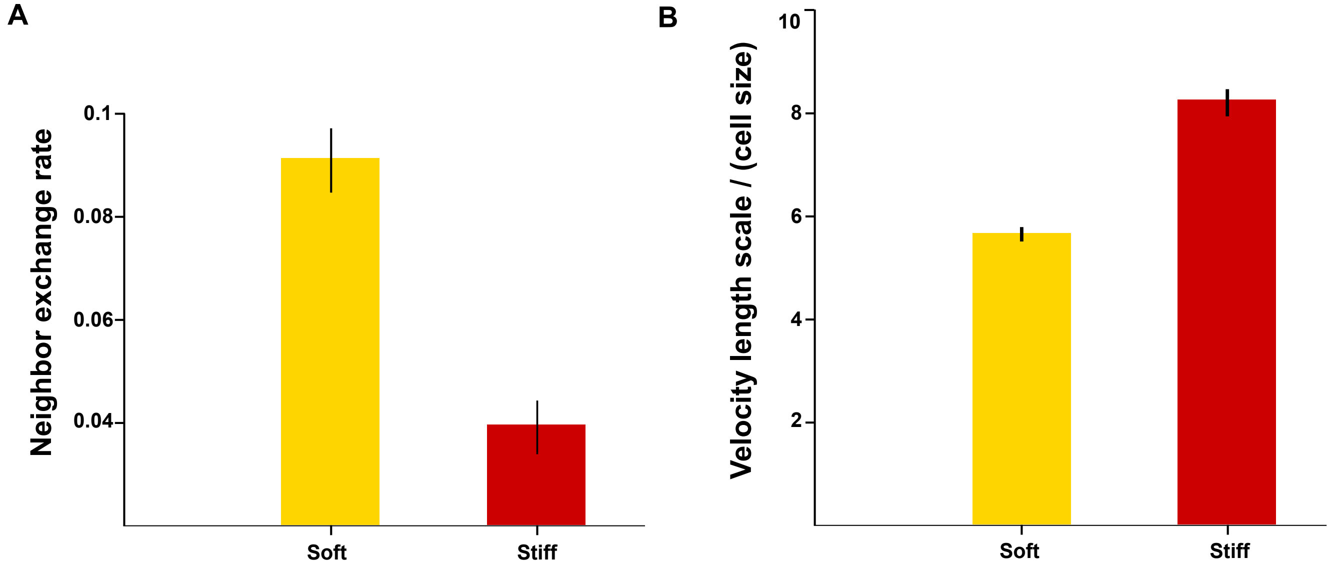


FIG. 6. **Fluid-like versus solid-like behavior of soft and stiff cells.** A) The rate of neighbor exchange in separate monolayers of soft ($\gamma = 0.025$) and stiff ($\gamma = 0.1$) cells. The rate is calculated per simulation time that corresponds to the full segregation time when soft and stiff cells are mixed. B) The velocity length scale normalized by the cell size for separate monolayers of soft ($\gamma = 0.025$) and stiff ($\gamma = 0.1$) cells. The length scale is calculated as the length over which the velocity-velocity correlation function $C_{vv} = \langle \boldsymbol{v}(r,0)\boldsymbol{v}(r,t)\rangle / \langle v(r,0)^2 \rangle$ in the monolayers crosses zero.

[20] I. S. Aranson, Physical Models of Cell Motility (Springer, 2016).
